# Cross species modeling reveals a role for the unfolded protein response in shaping the transcriptional reaction to *Mycobacterium tuberculosis* infection

**DOI:** 10.1101/2024.04.19.590290

**Authors:** Krista M. Pullen, Ryan Finethy, Seung-Hyun B. Ko, Charlotte J. Reames, Christopher M. Sassetti, Douglas A. Lauffenburger

**Author notes:** These authors contributed equally (Krista M. Pullen, Ryan Finethy). Co-corresponding authors Douglas A Lauffenburger, 77 Massachusetts Avenue, Cambridge MA 02139, **Email:**; Christopher M. Sassetti, 368 Plantation Street, Worcester, MA 01605, **Email:**. **Author Contributions:** Conceptualizing this research study: K.M.P, R.F., S.H.B.K., C.J.R., C.M.S., and D.A.L.; Analyzing data: K.M.P, R.F., and S.H.B.K.; Acquiring data: R.F., and C.J.R. Writing the manuscript: K.M.P, R.F., S.H.B.K., C.J.R., C.M.S., and D.A.L. Co-first authorship order was assigned in the order of initial involvement in the project. **Competing Interest Statement:** The authors have declared that no conflict of interest exists. Any opinion, findings, and conclusions or recommendations expressed in this material are those of the authors and do not necessarily reflect the views of the National Science Foundation or the Department of Defense. **Classification:** Biological Sciences, Immunology and Inflammation.

## Abstract

Numerous blood mRNA signatures have been developed to diagnose tuberculosis (TB) disease. The utility of these signatures in diverse populations depends on the inclusion of ubiquitously expressed features, such as type 1 interferon (IFN) production and innate immune cell activities. As a result, these signatures are generally insensitive to heterogeneous responses between individuals. Designing more effective therapies will require understanding the diverse mechanisms underlying pathogenesis by associating them with appropriate preclinical animal models. To address this critical animal-to-human gap, we applied a modeling framework, Translatable Components Regression, which is designed to account for biological heterogeneity by identifying multiple orthogonal axes of variation that are common to humans and animal models. Our framework was capable of distinguishing human active TB from latent TB infection using a model derived from murine data. This discrimination was based on differential expression of numerous biological pathways in addition to the common IFN and neutrophil signatures. Prominent among these predictive pathways was protein translation, which we show is a feature of the Mtb infection-induced Unfolded Protein Response (UPR) in macrophages. We show that this cellular stress pathway controls a variety of immune-related functions in Mtb-infected mouse macrophages, suggesting a possible causative role during the development of TB disease.

**Significance Statement:** Despite tuberculosis being one of the top causes of global mortality, the mechanisms that control the progression of disease are still not fully understand. Here we leverage a systems-level modeling approach that incorporates transcriptomics data across thousands of genes from both a traditional tuberculosis mouse model and human clinical samples to implicate a previously unappreciated mechanism in pathogenesis, the unfolded protein response. We validate these findings in a mouse macrophage model and pinpoint which branch of the unfolded protein response might be activated during tuberculosis infection. These insights, originally derived from our cross-species model, may allow us to better understand human tuberculosis pathogenesis and potentially identify therapeutic targets to prevent active tuberculosis.

## Introduction

Tuberculosis (TB) continues to be a major world-wide threat to human health as one of the leading causes of death from an infectious agent, second only to COVID-19 in 2022 (1). Exposure to the causative agent, *Mycobacterium tuberculosis* (Mtb), results in a spectrum of outcomes ranging from bacterial clearance to chronic asymptomatic infection and progressive disease of the lung or other anatomical sites (2). Despite this heterogeneity, Mtb infection outcome is often categorized in a binary fashion. Those that remain asymptomatic are classified as having latent TB infection (LTBI), whereas those with disease are considered to have active TB (ATB). This dichotomous framework is clinically useful but ignores the heterogeneity within each group which likely represents a variety of diverse and transient biological states (3). Genetic and epidemiological studies have revealed that Mtb infection outcome is influenced by genetic composition of both the host and pathogen, as well as a variety of nutritional and other environmental factors (4). This diversity in both risk factors and disease phenotypes implies heterogenous underlying mechanisms, which remain unclear.

To overcome the diagnostic challenges posed by the variable clinical presentation of tuberculosis (TB), the past decade has seen a major effort to leverage blood mRNA signatures to detect disease (5). Dozens of signatures have been reported to differentiate ATB from LTBI and other infections, to identify those with incipient disease, or to quantify treatment efficacy (6–11). These signatures are commonly composed of genes corresponding to the type I IFN response and neutrophils, and they are effective in detecting and/or quantifying TB disease. Despite these advances, the vast majority of human transcriptomic studies in tuberculosis have focused on diagnosis, and therefore only discovered common features of TB disease that are shared broadly between individuals. This type of analysis explicitly excludes heterogeneity between individuals that likely reflects important mechanistic differences in immunity and pathogenesis. Indeed, a meta-analysis of 45 published tuberculosis gene signatures that mostly include less than 100 genes, found 1513 unique genes to distinguish between TB phenotypes in different settings (12). This suggests high prevalence of cohort- and phenotype-related heterogeneity, potentially reflecting diverse underlying disease mechanisms. Similarly, a previous modular analysis of blood mRNA data from Mtb infected humans identified a number of independently-varying immune features and pathways that differentiate the ATB from LTBI phenotypes (13). Understanding the diversity of biological processes that underlying TB disease progression is crucial for the development of broadly effective interventions, such as vaccines or immunotherapies.

A further challenge in therapeutic development is relating diverse human phenotypes with the more homogenous animal models where mechanistic and preclinical studies can be performed (14). Historically, inbred mouse strains were extensively used as a murine model for ATB, but generally lack the heterogeneous outcomes seen in humans. Genetically diverse mice (15, 16), as well as non-human primates (17), have been developed as animal models that more faithfully reflect the diversity observed in natural populations. In each case, transcriptomic features that differentiate ATB disease states across species can be identified (18, 19). These observations suggest shared disease biology between species, which bolsters translationally-relevant mechanistic studies in the small animal model that suggest causative roles for these pathways. Nonetheless, fundamental immunological differences exist between species (20, 21), many of which are relevant to TB disease, highlighting the continued challenge of translating biological insights between animal models and humans.

Traditionally, cross-species analyses comprise direct, one-to-one comparisons of the presence, or absence, of particular biomolecules (genes, proteins, and/or metabolites) or cell types. The goal of these studies is often to find dominant biomarkers that are shared across all individuals of both species. As a result, these comparisons typically exclude heterogeneous disease mechanisms as well as those that are subdominant in one of the species. Toward addressing this problematic preclinical-to-clinical gap, Brubaker et al. developed a novel modeling framework known as Translatable Components Regression (TransComp-R) that is designed to account for biological heterogeneity by identifying multiple orthogonal axes of variation in one species that are correlative with disease biology and phenotypes observed in another species (22). Applying this framework in the context of inflammatory bowel disease (IBD), the study demonstrated that mouse proteomic features could provide insights relating to human transcriptomic expression, allowing for better classification of patient responses to IBD treatment and identification of novel pathways implicated in therapy resistance. TransComp-R has since been applied, and extended to translate gene pathway analyses, with success in the realm of neuropathologies but has yet to be applied in the context of infectious disease (23, 24).

Here, we adapt TransComp-R to identify biological pathways distinguishing human LTBI from ATB based on data from a murine TB model, using data from a study by Moreira-Teixeira et al. (25). This study illustrated that a transcriptional signature prominently found in humans with ATB, reported by Singhania et al. (26), could also be found in a cohort of mice designed to reflect different degrees of disease via the inclusion of two mouse strains, with differing susceptibility to infection, and two Mtb isolates administered at different doses. Instead of defining diagnostic signatures, our approach is designed to maximize the amount of information we can acquire about heterogeneous ATB and LTBI disease mechanisms from mouse models, both gaining insights about the relationship between mouse and human TB pathology and ultimately utilizing mouse data to further our mechanistic understanding of human infection. We demonstrate that a mathematical model built on the mouse blood transcriptomic data accurately predicts human disease state. This model is composed of multiple orthogonal axes of variation that individually correlate with human outcome and represent biologically distinct pathways. We identified a diverse range of associated pathways, in addition to the commonly identified IFN and neutrophil signatures. Prominent among these predictive pathways is protein translation, which we show is a feature of the Mtb infection-induced Unfolded Protein Response (UPR) in macrophages. We demonstrate that this cellular stress pathway controls a variety of immune-related functions in Mtb-infected mouse macrophages, suggesting a possible causative role during the development of TB disease. Overall, we illustrate how translational cross-species modeling allows us to hypothesize previously unappreciated biological mechanisms driving TB disease and identify potential therapeutic targets.

## Results

### Direct comparison of gene expression across species reveals molecular discrepancies between preclinical models and human TB disease

Since the aim of this study is to learn more about the relationship between the transcriptional response in tuberculosis animal models and human tuberculosis patients, we utilized mouse and human bulk blood transcriptomics datasets previously published in studies by Moreira-Teixeira et al. (25) and Singhania et al. (26), respectively. The murine tuberculosis model used in this study contrasted C57BL/6J with the relatively sensitive C3HeB/FeJ mouse strain to reflect differing degrees of disease. In addition to utilizing both mouse strains, Moreira-Teixeira et al. incorporated two different Mtb strains, given at two different doses to different cohorts of mice (**Figure 1A**). Due to the excessive pathology in some groups, the blood sample for transcriptomic analysis was collected at different time points for different mice, always at least 26 days after infection with Mtb. As was done by Moreira-Teixeira et al, we compared this murine cohort to human transcriptomic data from the Singhania et al. study, focusing our study on the London and South Africa cohorts (**Figure 1B**). Typically, cross-species analyses of molecular data involve direct, one-to-one comparison of abundance or expression. As such, we compared the median expression of the twenty gene signature identified in humans by Singhania et al. to the mouse cohort (**Figure 1C**). We observed that 25% of genes that were more highly expressed in a particular human phenotype switched to be higher in the opposing phenotype in mice. In other words, genes that were more highly expressed in human ATB had relatively higher gene expression in C57BL/6J mice that develop relatively less severe disease, and vice versa. Additionally, we could not identify a mouse homolog for one of the twenty human genes, APOL4. When comparing the magnitude of the median gene expression, the range in gene expression was larger between active and latent human TB patients than between C57BL/6J and C3HeB/FeJ mice (**Figure S1**). To assure that our observation of a less prominent, and sometimes inconsistent, TB gene signature in mice was not specific to the Singhania et al. gene set, we compared the enrichment of 45 different published TB gene sets in each of our phenotypes by species (**Figure 1D**). While these signatures distinguished human phenotypes, we observed a similar trend of weak differentiation of mouse or Mtb strains, highlighting a gap in our ability to translate between species solely using direct comparison of the expression of homologous genes.

**Figure 1.**
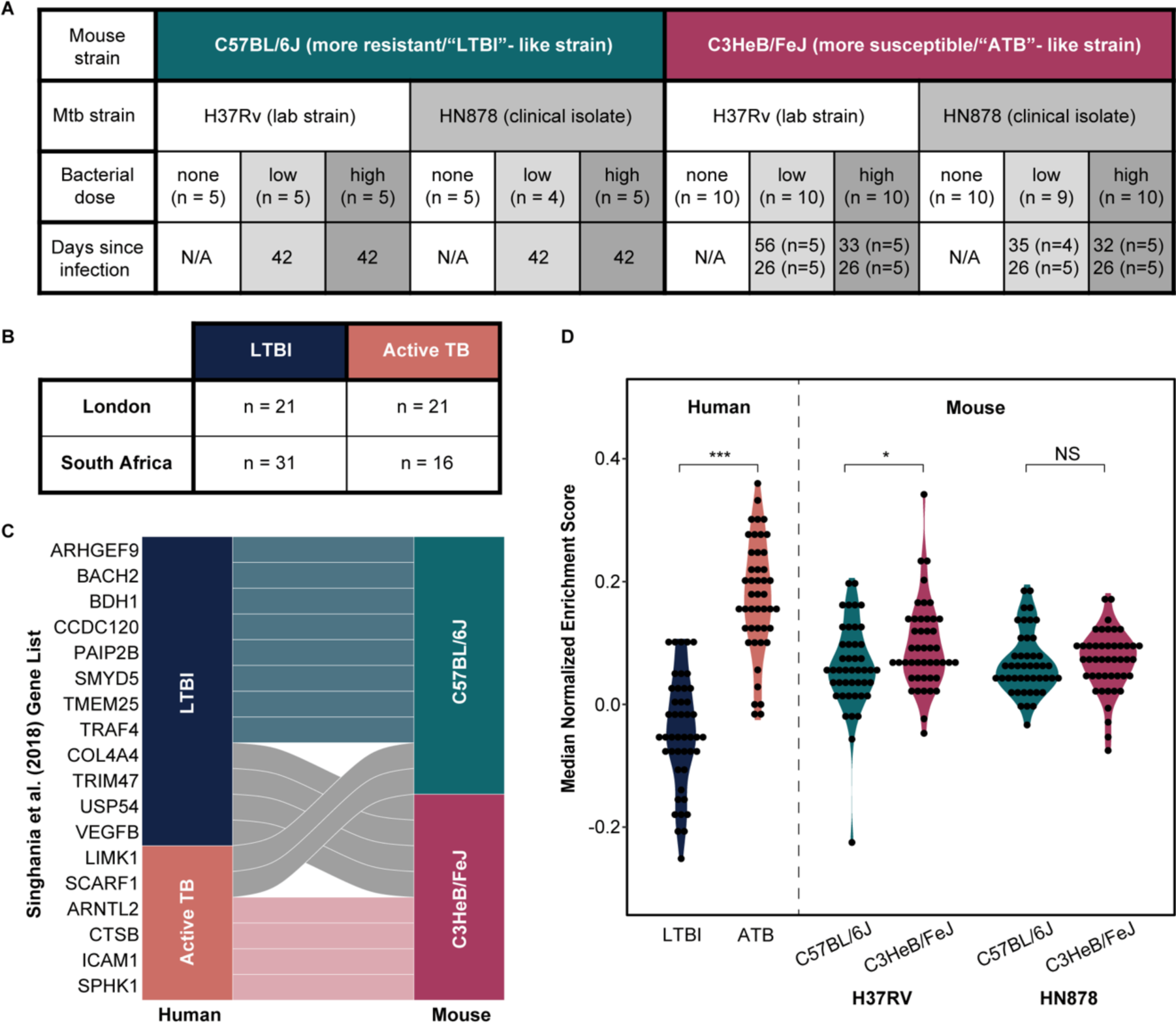
Direct Comparison of Gene Expression Across Species. (A) Table summarizing the mouse model. (B) Table summarizing the human tuberculosis study. (C) Alluvial plot illustrating relative gene expression of the published Singhania et al. (2018) human gene set in human active vs. latent tb and the two mouse strains. Each gene is classified as more highly expressed in one of the two phenotypes in each species and the lines in between each bar illustrate whether the gene is higher in the latent cases of each species (blue), active case (pink) or is inconsistent between species (gray). APOL4 is not shown because there is no known mouse homolog. KCNC4 is not shown because the median expression level in both mouse models was equivalent. (D) Single sample GSEA where each gene set is one of 45 published TB gene signatures, containing at least 10 genes, for discriminating between active and latent disease. The median enrichment of the gene set in each phenotype (i.e.-active or latent) is plotted as one data point.

### TransComp-R model identifies murine transcriptomics signatures relevant to human TB disease phenotypes

Rather than comparing individual genes on a one-to-one basis between species, we sought to construct a cross-species model that identifies transcriptomics-derived axes of variation in the mouse tuberculosis model that correlate with human tuberculosis phenotypes and may allow us to discover novel mechanisms driving disease that are typically overshadowed by IFN and neutrophil signatures.. To minimize random noise, we chose to build the foundations of our model on genes that were differentially expressed (DEGs) in our human cohorts, and their mouse homologs (**Figure S2**). 3,201 direct DEG homologs between the mouse and human datasets were identified for inclusion in our cross-species model.

The TransComp-R framework utilizes both human and animal data to develop a better understanding of the relationship between animal models and human biological mechanisms, ultimately allowing us to develop further understanding of the human disease pathology, progression, and/or prevention. Briefly, dimensionality reduction is first applied to the mouse model to generate a small number of latent variables, here also known as principal components, that capture the majority of the variation in the mouse data (**Figure 2A**). Then, using the feature (e.g., gene) loadings for each mouse principal component (mPC), the human homologs are projected into the mouse latent variable space. In other words, the human data is transformed based on the relative importance of each mouse homolog such that the human data can be described in terms of distribution across mPCs. Since there is no guarantee that mPCs that describe the most variation in the mouse data will capture differences in the human phenotype of interest (e.g., LBTI vs. ATB), the mPCs with the greatest univariate effect size between LBTI and ATB samples were selected to be included in the final logistic regression model. Regressing the human phenotypic classifications onto the selected mPCs validates the importance of the selected mPCs, and thus the relative importance of each mouse gene, in discerning between human disease states. From here, gene set enrichment analyses can be performed to garner understanding of the biological pathways in mice that correlate with active or latent tuberculosis in humans.

**Figure 2.**
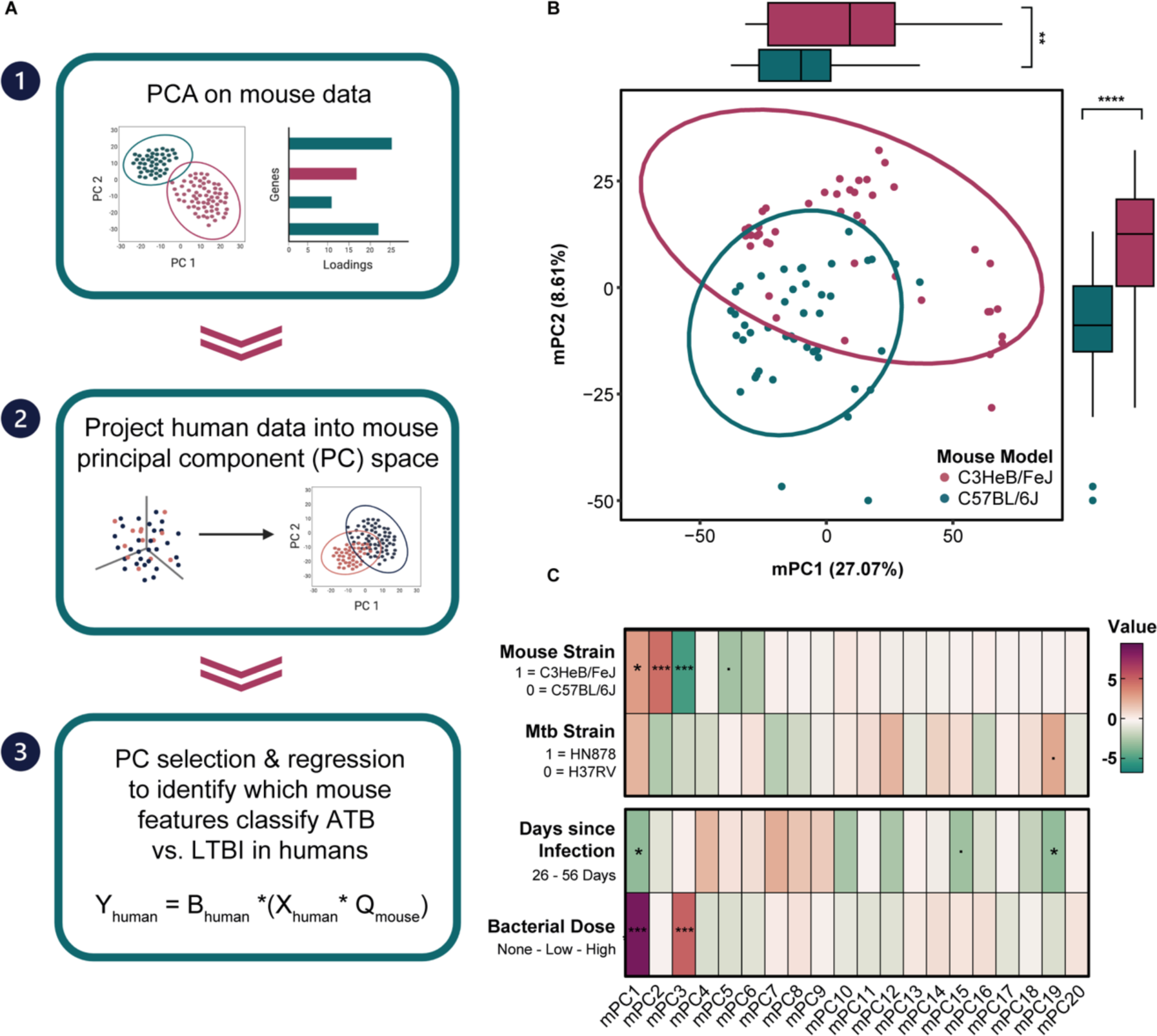
TransComp-R model constructed from mouse transcriptomics data. (A) Schematic of the TransComp-R workflow. (B) Principal component analysis on mouse homologs of the human differentially expressed genes in latent vs. active tuberculosis. Each data point represents a mouse sample, and the colored ellipses represent the 95% confidence interval for each strain’s distribution in the PC1 vs. PC2 space. Boxplots on the x-and y-axes illustrate the distribution of the C3HeBFeJ (pink) and C57Bl6 (blue) mouse strain samples on each axis, respectively. Wilcoxon rank sum tests were performed on the scores of each mouse strain for each principal component to determine if the univariate distributions were statistically significantly different. ** p-value < 0.01; **** p-value < 0.0001(C) Regression model results from individual mouse principal components regressed onto mouse strain, Mtb strain, days post-infection, and bacterial dose. The Mtb strain and days since infection models do not include uninfected controls. Z-values are shown for the logistic regression models (mouse and Mtb strain). T-values are shown for the linear regression models (days post-infection and bacterial dose). Positive values (magenta) correspond with the C3HeBFeJ mouse strain, HN878 Mtb strain, longer time post-infection and higher bacterial dose. ‘.’ indicates p-value < 0.1, * indicates p-value < 0.05, and *** indicates p-value < 0.001 after Benjamini-Hochberg multiple hypothesis correction.

As detailed above, we built the foundations of our cross-species model by performing principal component analysis (PCA) on the 3,201 DEG homologs in the different mouse cohorts (**Figure 2B**). Separation between the two mouse strains is noticeable on the first two mPCs, with statistically significant univariate distribution between strains along each mPC. This did not come as a surprise based on existing literature on the distinct susceptibility and disease pathologies observed in C57BL/6 and C3HeB/FeJ mice. As illustrated in **Figure 1A**, the mouse data had several known sources of variation beyond mouse strain. Explicitly, *Mtb* strain, *Mtb* dose and days post-infection for sample acquisition also varied between samples. To distinguish which, if any, mPCs these variables influenced, regression models were built to regress each covariate onto the scores of each mPC, individually (**Figure 2C**). This analysis revealed that most of these sources of variation, excluding Mtb strain, do significantly correlate with the distribution of scores on various principal components, especially mPC1. We can also see this visually when we color the scores plot by each covariate (**Figure S3**). Each of these sources of variability (mouse strain, Mtb strain, dose, and time point) could reflect different aspects of human disease, we continued our study with all the mouse data to include the greatest possible extent of molecular variation in the data structure of our model, with hopes of most effectively translating these highly controlled mouse data to naturally heterogenous human populations. The first 20 mPCs explain 75% of the variance across the murine data and, thus, were included in the next step of the model (**Figure S3**).

Next, data from the human TB cohorts were projected into our mouse principal component space, resulting in human sample scores along the mPC axes. The projected samples showed substantial separation by TB disease status (LTBI versus ATB) on mPC1 vs mPC2, with 45% of the total variance captured by these two mPCs (**Figure 3A**). This indicates that the mouse molecular data structure possesses information relevant to distinguishing between human TB phenotypes across multiple mPCs. The contributions to mouse data variance decrease as the order of the mPC increases, as expected, while the contributions to human data variance do not necessarily follow the same order monotonically (**Figure 3B**). This suggests that common biological pathways explain different degrees of variation in the two species. Consequently, while 20 mPCs are required to explain 75% of the variation in the mouse transcriptional response to Mtb, 75% of the variation in the human transcriptomics is described over just 7 mPCs (**Figures 3B and S4A**). Thus, only the first 7 mPCs were included in the following steps of the model. The only human covariate, other than disease phenotype, that appeared to significantly correlate with any of the selected mPCs was the patient’s gender as reported in the study by Berry et al. in which the original clinical samples were collected (**Figure S4B**). For comparison, PCA was also performed on the human data directly (**Supplementary Figure S5A**). Interestingly, while significant separation between ATB and LTBI was observed on the first human PC, the distribution of samples on the second human principal component was not statistically significant. In other words, there are more distinct axes of transcriptional variation in the mouse model that correlate with distinction of human ATB and LTBI than there are in the model constructed directly from the human transcriptomics data. It is plausible this phenomenon occurred because human PC1 alone can describe the majority of the disease status-specific variation as the model was built on human DEGs with significant differences between ATB and LTBI phenotypes. Alternatively, in the human data, the signal from genes involved in pathways classically activated by Mtb infection, such as IFN response, may shroud more subtle signals relating to pathways only activated in a subset of human TB patients.

**Figure 3.**
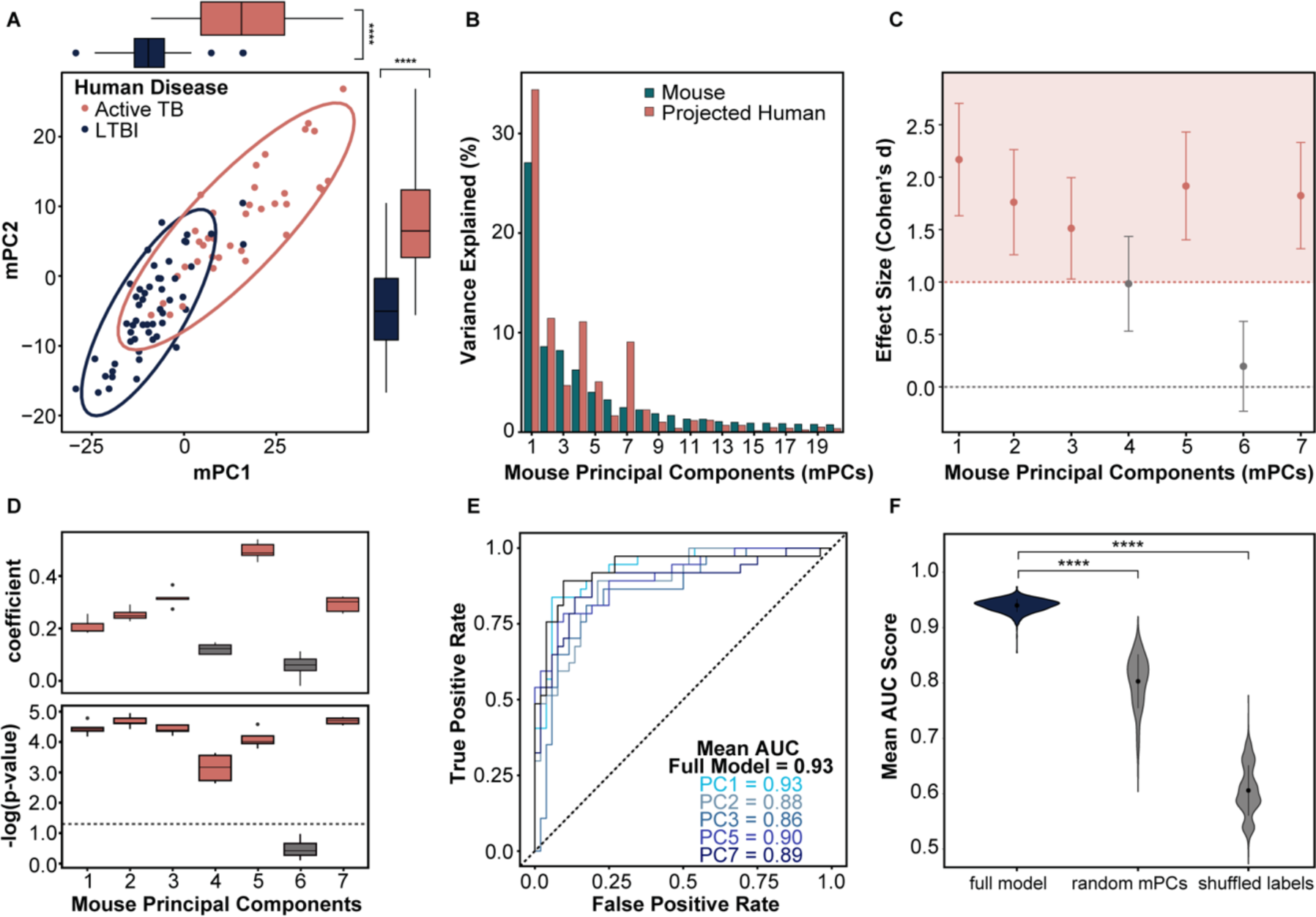
Murine-derived principal components distinguish human TB phenotypes. (A) Human latent vs. active tuberculosis samples projected into the mouse PC1 and PC2 latent variable space. Each data point represents a human sample, and the colored ellipses represent the 95% confidence interval for each disease group’s distribution in the PC1 vs. PC2 space. Boxplots on the x- and y-axes illustrate the distribution of the active tuberculosis (salmon) and LTBI (navy) samples on each axis, respectively. Wilcoxon rank sum tests were performed on the scores of each human phenotype for each principal component to determine if the univariate distributions were statistically significantly different. (B) Percent variance explained by each mouse principal component for the human and mouse data. (C) Plot comparing the univariate effect size observed between the ATB and LTBI human samples projected into each individual mPC. Effect sizes, measured by Cohen’s d coefficient, with a magnitude greater than one are shaded pink. Confidence intervals for each Cohen’s d coefficient are indicated by error bars. (D) The top boxplot shows the regression coefficient for each mPC when individually regressed on human outcome (ATB vs. LTBI). The bottom plot illustrates the corresponding p-values, adjusted with Benjamini-Hochberg multiple hypothesis correction. The dotted line indicates a p-value of 0.05. In both plots, the whiskers represent the IQR across 5-folds of cross validation (CV). (E) Mean ROC curves for 100 trials of 5-fold logistic regression of human disease phenotypes using projected human data onto mPC1, mPC2, mPC3, mPC5, and mPC7 individually and together (full model). (F) Mean area under the ROC curve (AUC) of 100 trials of 5-fold CV on the full TransCompR model compared to null models with five randomly selected mPCs and null models with mPC1, mPC2, mPC3, mPC5, and mPC7 with shuffled phenotype labels. P-values were calculated from the tail probability of the null distributions.

Toward our goal of finding biological insights in the mouse data pertinent to human disease presentation, we then used the distribution of scores from the projected human data to calculate univariate effect size between phenotypes on each mPC individually (**Figure 3C**). From this analysis, the magnitude of effect size along 5 mPCs stood out among the first 7 mPCs: mPC1, mPC2, mPC3, mPC5, and mPC7. When ATB and LBTI phenotypes were regressed onto these selected mPCs individually, we observed similar predictive capabilities (**Figure 3D-E**). While regression of mPC4 had statistical significance (p-value < 0.05), we excluded it from subsequent analyses as both the effect size and regression coefficient of mPC4 was lower than the other 5 statistically significant mPCs. The mouse principal component with the largest magnitude regression coefficient was not mPC1, but rather mPC5, highlighting the importance of including multiple mPCs in our analysis. The top five mPCs (mPC1, mPC2, mPC3, mPC5, and mPC7) were capable of predicting human ATB versus LTBI phenotype labels with high classification accuracy in logistic regression models of individual mPCs and together in one full model (**Figure 3E**). Furthermore, the full model had significantly higher accuracy than null models with random mPCs or shuffled phenotype labels. These results suggest that our identification of five axes of variation (mPCs) in the mouse model that significantly correlate with human ATB and LBTI disease phenotypes was not due to random chance (**Figure 3F**). Furthermore, while mPC1 may possess the most information relevant to variation in the mouse TB model, the other four mPCs appear to contain distinct yet pertinent information relevant to differentiating human TB phenotypes as well. Based on the proportion of human variance explained by each axis and their subsequent performance in classifying human TB disease states, mPC1, mPC2, mPC3, mPC5, and mPC7 were deemed to be the “Translatable Components” (TCs) of this model.

### Translatable components represent biological axes both similar to and distinct from known TB-associated mRNA signatures

To evaluate the similarity between the TCs identified from our TransComp-R model and previously identified biomarkers of TB, we compared the gene loadings on each principal component with 67 published blood mRNA signatures of TB. These signatures varied in size from 10 to 893 genes and were derived from both demographically distinct cohorts and meta analyses (12). Using a single sample GSEA scoring metric, we found that the majority of these signatures were most highly enriched on mPC1 (**Figure S6**). Signature scores for TC1 and TC3 were significantly higher than each of the other TCs which were generally not significantly different from each other, with exception to TC5 and TC7. This observation is consistent with the greatest amount of variance in human gene expression being explained by TC1 (**Figure 3B**). It is notable that these include signatures based on relatively distinct cohorts or analytical methods. For example, this group includes signatures of very early progression (27) or pediatric TB (28). It also includes one based on a modular transcriptional analysis designed to detect heterogeneous responses (26). Thus, we conclude that TC1 and TC3 capture commonly described TB signatures, while the other TCs might contain novel heterogeneous biology that is typically masked by the dominant TC1-related response and/or is more characteristic of TB disease states other than the common pulmonary TB versus LTBI contrast.

### Translatable Components reveal diverse immunological pathways

Due to the orthogonal nature of TCs, we expected that each would reflect distinct cellular processes. To investigate this potential heterogeneity, we used Ingenuity Pathway Analysis (IPA) to identify the functional pathways most associated with each TC, and subsequently examined the association of each pathway with each TC **(Figure 4A, Table S1)**. As anticipated, each TC associates with unique combinations of functional pathways. Consistent with its similarity to known TB-related mRNA signatures, TC1 strongly associates with established disease-related pathways, specifically those related to neutrophil activity and interferon signaling. TC2 gene loadings demonstrated comparable alignment with interferon related gene sets but was not associated with neutrophil signaling. Notably, the gene loadings for the later TCs (TC2, TC3, TC5 and TC7) all reflect significant, though variable, associations with translation pathways, alongside different immunological pathways that define their unique identities, e.g. mPC3 also aligns with interferon gamma signaling and natural killer cell activity.

**Figure 4.**
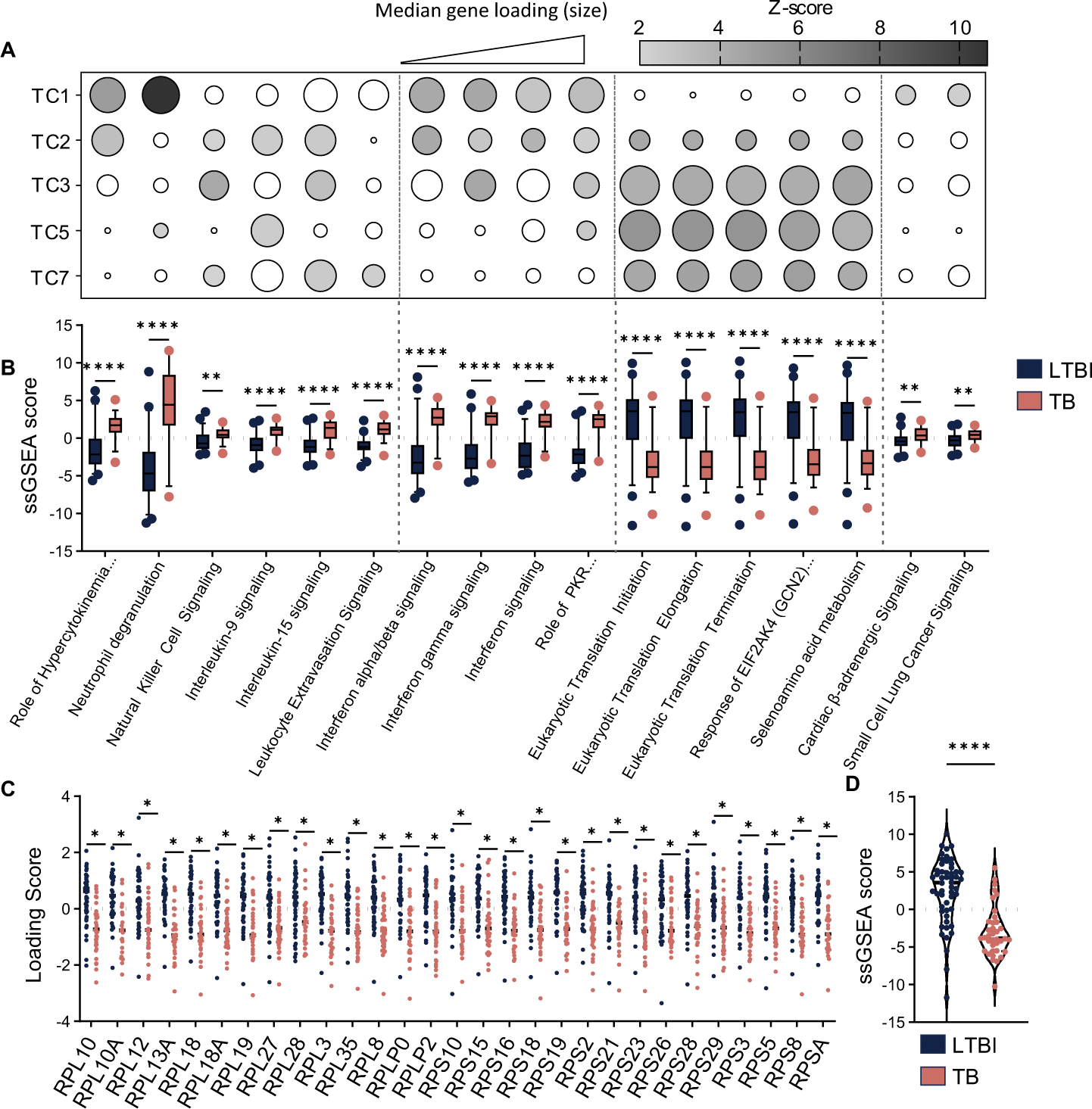
Translatable components reflect diverse biological functions including protein translation. Pathways most associated with individual TCs were identified using IPA and categorized by function. (A) Dot plot heatmap illustrating association of all TCs with selected pathways. Shading represents absolute value of z-score, with unshaded circles indicating non-significant z-score. Dot size corresponds to the absolute value of the median loading score for leading-edge genes. To align gene expression data with pathway activity, the sign of inversely correlated genes was reversed prior to calculating the median. (B) Comparison of ssGSEA signature scores for each selected pathway between patients categorized as LTBI or ATB. Box plots depict median and 5-95 percentiles. Statistical analysis was performed using multiple unpaired t-tests with Holm-Šídák multiple comparisons test. Leading-edge genes across the translation-associated pathways were compared to generate a common gene set. (C) Comparison of loading scores for individual translation genes within the human cohort. Significance was determined by multiple unpaired t-tests with Holm-Šídák multiple comparison correction. (D) Composite analysis of the translation gene set using ssGSEA across the human cohort. Statistical analysis was performed using unpaired t-test. * p-value < 0.05, ** p-value < 0.01, **** p-value < 0.0001.

To affirm the disease-relevance of these pathways, we evaluated the strength of association between TC-aligned pathways and human TB disease state. Using ssGSEA, signature scores were calculated for each individual in the human cohort and compared between patients categorized as LTBI or ATB **(Figure 4B).** We found that each pathway was differentially enriched in these two groups. As expected, signature scores for established disease pathways, e.g. those relating to neutrophils or interferon signaling, effectively differentiate between latent and active TB. Translation pathways were found to exhibit comparable differential enrichment between these groups.

Our analyses suggested that altered expression of translation-related genes is an important translatable feature of TB disease. To better understand this association, we compared the leading-edge genes across the translation pathways to identify common elements, revealing a shared gene set that is comprised of ribosomal components. For the shared gene set, we compared the expression of the individual genes across the human cohort **(Figure 4C),** finding that each of the 29 genes was relatively repressed in the ATB group. Composite analysis of the gene set using ssGSEA similarly revealed significant differential expression between LTBI and ATB groups **(Figure 4D)**. These results reveal that ribosomal gene expression is predictive of disease.

### Ribosomal mRNA repression is a feature of Mtb infection of macrophages and depends on the unfolded protein response (UPR)

Translational inhibition is a hallmark of the UPR, a cellular stress response triggered by proteotoxic stress in the endoplasmic reticulum or mitochondrion. The response can be induced upon infection with intracellular bacterial pathogens, and markers of UPR activation have been reported in human TB lesions. Based on these observations, we hypothesized the regulation of ribosomal mRNAs associated with the TCs may be linked to UPR activity in infected cells. UPR signaling is propagated by three distinct pathways initiated by the inositol-requiring enzyme 1 (IRE1α), activating transcription factor 6 (ATF6) or protein kinase R-like endoplasmic reticulum kinase (PERK). Each of these pathways contributes to the ultimate effects of UPR activation, which can include induction of chaperones, translational inhibition, and the potentiation of pro-inflammatory signaling. To understand the influence of the UPR during Mtb infection, we assessed the impact of inhibition of the three canonical UPR pathways on the macrophage transcriptional response to the pathogen.

Mouse BMDMs were infected with Mtb in the presence or absence of chemical inhibitors targeting IRE1α (4μ8c), ATF6 (Ceapin-A7), and PERK (GSK2656157) and mRNAs were quantified by RNAseq. We confirmed the inhibitors to be efficacious in BMDMs by assessing mRNA levels of known targets during tunicamycin-induced ER stress, and found that these compounds did not affect the intracellular uptake of Mtb across a range of concentrations **(Figure S7A-B)**. Cluster analysis of the transcriptomic results revealed distinct expression patterns corresponding to the inhibition of each UPR pathway during infection **(Figure 5A, Table S2)**. Using IPA, we found that each cluster is associated with pathways corresponding to different cellular functions. These clusters were primarily related to gene sets representing the following functions: cluster 1 – cell cycle, cluster 2 – interferon and other cytokine production/signaling, cluster 3 – translation, 4 – various signaling pathways, 5 - ubiquitination and 6 - extracellular matrix. To assess the impact of infection and UPR inhibition on the pathways identified using TransComp-R, we used IPA to calculate enrichment scores for the major TC-associated pathways **(Figure 5B, Table S3)** in the BMDM data. Infection of BMDMs produced a transcriptional response that includes many of the human TB-associated pathways identified in our TC analysis, e.g. interferon signaling and translation pathways. UPR inhibition limited the infection-induced changes in expression of these same pathways to different extents. All three inhibitors influenced pathways related to cytokine production, although inhibition of PERK exhibited a more muted effect. Inhibition of ATF6 and IRE1α, but not PERK, limited interferon signaling. Translation associated pathways were only significantly associated with ATF6 inhibition. These effects inferred from pathway enrichment scores were also evident by the differential expression of leading-edge genes from the TC analysis in three of the major pathways. In particular, ATF6 inhibition completely reversed the Mtb infection-induced repression of the consensus protein translation gene set. **(Figure 5C)**. These data suggest that significant aspects of the cross-species translatable features of Mtb infection could be attributable to UPR activation.

**Figure 5.**
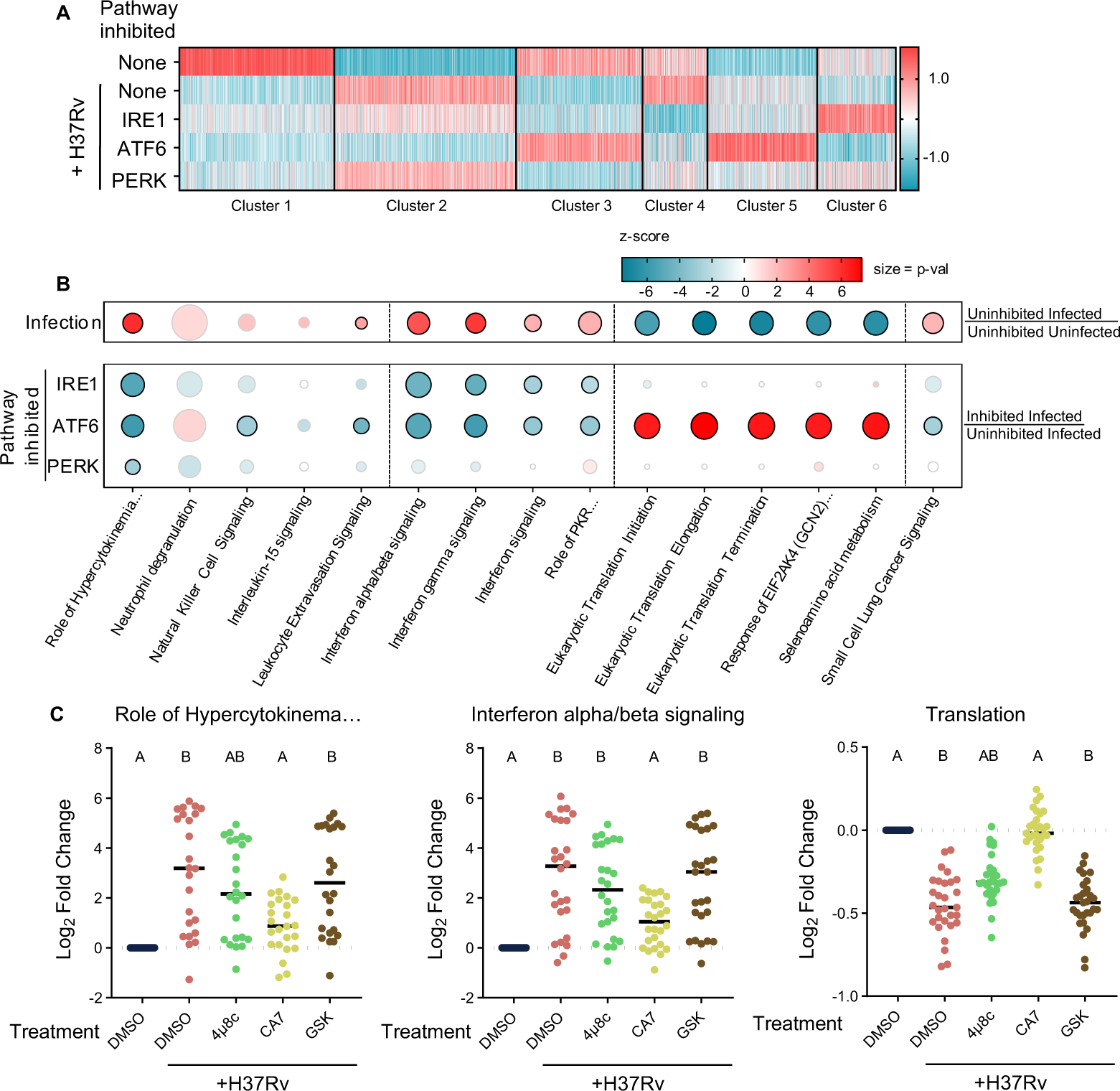
UPR activation influences multiple TB-associated pathways in BMDM and ATF6 is particularly associated with protein translation. BMDMs were infected with H37Rv in the presence or absence of UPR inhibition or left uninfected. Following infection, mRNAs were quantified via RNAseq. Normalized counts were averaged and K-means clustering was applied to group genes based on differential expression patterns. (A) Heatmap showing results of clustering. (B) Dot plot heatmap showing associations with Mtb infection (top) and UPR inhibition during infection (bottom). Color is indicative of z-score while size represents p-value. Pathways with significant enrichment are outlined in black. Major-TC pathways with no significant association (p-value) under any condition were excluded. (C) Expression of leading-edge genes from TC analysis for indicated pathways. Log2fold change relative uninfected BMDMs is shown. To align gene expression data with pathway activity, the sign of genes whose expression was inversely correlated with pathway activity were reversed. Statistical comparisons were made using nonparametric Friedman test with Dunn’s multiple comparison’s test. A = p-value < 0.05 when compared to infected uninhibited condition. B = p-value < 0.05 when compared to uninfected condition.

## Discussion

The diversity in both risk factors for human TB disease progression and the phenotypes observed following Mtb infection suggest that numerous underlying biological processes likely influence the trajectory of disease. This heterogeneity is a challenge to the development of broadly effective interventions (3). Primary among these challenges is the complex relationship between the heterogeneous human TB pathophysiology and the more homogenous small animal models used for discovery and preclinical studies (14, 29). Indeed, the assumption that animal model biology translates directly to human disease has led to a number of failures in TB therapeutic development, as well for many other human diseases (30, 31). Our lab has developed a cross-species modeling framework, TransComp-R, to overcome animal model limitations by identifying axes of molecular variation in preclinical models that correlate with the observed human disease pathologies, while acknowledging that each individual pathway may be variably important between individuals or species (22). The application of this translational modeling framework in other disease contexts identified novel disease-relevant biological pathways, which were not apparent when only the commonly dominant features across both species were considered (22–24). Here we apply TransComp-R to TB in efforts to untangle the heterogeneity underlying human TB disease presentation, and similarly identify multiple diverse disease-associated biological axes that are shared by humans and mice. Strikingly, the relative importance of each axis of transcriptional variation is distinct between species. This study reveals that a previously unappreciated degree of heterogeneity relevant to human TB is present in a traditional mouse model of Mtb infection and implicates new biological pathways, such as the UPR, underlying distinct human TB phenotypes.

Similar to gene expression signatures defined directly in human cohorts, our mouse PC model effectively discriminated human LTBI from ATB. By projecting human clinical data into our murine transcriptomics model, we were able to identify biological pathways enriched in mice that correlate with human phenotypes, but can be overshadowed by more canonical molecular responses, such as type I IFNs. While our model still captured previously published biological mechanisms in response to Mtb infection, it also provided additional insight into the relationship between the two species and identified novel pathways potentially involved in human TB pathogenesis. Existing TB signatures were most similar to TC1 of our model, which explains approximately a third of the variance in both human and mouse datasets. TC1 was also predominantly associated with the bacterial dose that the mouse was exposed to, suggesting that the gene expression features corresponding to both TC1 and many existing signatures represent a quantitatively dominant response to bacterial burden. This inference is supported by the sensitivity of TB diagnostic gene expression signatures to antimicrobial therapy (32).

In contrast to the reductive approaches used to derive diagnostic mRNA signatures, TransComp-R is designed to identify both dominant and subdominant features reflective of heterogeneous biology. Previous work using a modular gene expression approach supported the presence of heterogeneously expressed genes in the same dataset used in our study (25, 26). The model we present quantifies the importance of these subdominant features, identifying 4 mouse-derived PCs that together explain approximately the same amount of variance in the human dataset as PC1. These axes serve to connect distinct infection-related stimuli with corresponding gene expression responses. Whereas TC1 was associated with bacterial burden and canonical TB-related pathways, TC2, TC5, and TC7 were associated with the mouse strain and/or Mtb strain differences and reflected distinct immunological pathways such as NK cells and IL9 signaling. In sum, these observations support a conceptual model in which heterogeneity in gene expression can be assigned to functionally orthogonal axes, which are predominantly influenced by either antigenic burden or genetic variation in the host or pathogen. Importantly, each of these distinct axes is individually predictive of TB disease state.

Prominent among the pathways associated with the subdominant TCs (TC2, TC3, TC5, and TC7) were those related to protein translation. In both human blood and mouse BMDM the corresponding genes were repressed by ATB or Mtb infection, respectively, and this response depended upon the UPR. The ATF6 branch of the UPR was primarily involved in this response, with a more modest contribution of IRE1α. This cross-species translatable response to Mtb infection supports a likely role for UPR activation in both situations. Previous work demonstrating UPR-related gene expression in human TB lesions further supports this model (33). We acknowledge that the anatomical and cellular context was different in our mouse and human datasets, and direct infection of BMDM does not fully recapitulate the dynamics of human blood where diverse cell types are present and viable Mtb is quite rare. As a result, we speculate that UPR activation is not only a feature of directly infected macrophages but is also more generally related to immune stimulation. UPR activation has been observed in response to cell stimulation by diverse ligands, including TLR2 agonists (34), and systemic administration of inflammatory cytokines can produce UPR activation in tissues (35). While the specific cell(s) responsible for UPR-related gene expression in human blood remain unclear, our data suggest a mechanistic basis for an important component of diagnostic signatures that discriminate LTBI from ATB.

Inhibition of translation is a canonical function of the UPR, though transcriptional repression of ribosomal gene expression has not been described in mechanistic detail. Our data suggest that ATF6 is necessary for this effect, though we cannot differentiate between direct and indirect effects of this transcription factor. In addition to the canonical effects on translation and chaperone function, UPR activation has been implicated in a wide variety of cellular immune responses, including NF-κB activation, type I interferon production, and antigen presentation (34, 36–39). Our BMDM data support an important role for the UPR in shaping the myeloid transcriptional response to Mtb infection and demonstrate that the different branches of the UPR have distinct effects. ATF6 has the strongest effect on translation, IRE1a and ATF6 both repress a cluster of genes related to IFN and other cytokines, and these two branches have opposite effects on ECM-related genes. While we found PERK to have a relatively modest effect in C57BL/6 BMDM, previous work implicated this branch of the UPR in disease progression in C3HeB/FeJ mice – the same strain included in our TransComp-R study. Together these data suggest an important role for UPR activation, not only generating diagnostic mRNA signatures, but also in shaping the immune response to Mtb infection. While follow-up studies will be necessary to determine whether molecular components of the UPR could as a potential TB therapeutic target, this study highlights the utility of translational cross-species modeling as it has the potential to increase the value of preclinical data and expand our understanding of the human immune response to Mtb infection and accelerate the development of effective interventions.

## Materials and Methods

### Cohorts

As in the Moreira-Teixeira et al. study (25), we made use of human transcriptomic datasets from the study by Singhania et al. (26), which comprised cohorts from London ( n=21 LTBI, n=21 ATB), and South Africa (n=31 LTB, n=16 ATB). We excluded the Leicester tuberculosis progressor cohort from our analysis as progression between disease phenotypes was outside of the scope of our study. We aggregated these into one large human cohort with total n=89. We also utilized the mouse dataset from Moreira-Teixeira et al. comprised of C57BL/6 mice infected with either the H37Rv laboratory *Mtb* strain (n=5 uninfected, n=5 low-dose infected, n=5 high-dose infected) or the HN878 clinical *Mtb* isolate (n=10 uninfected, n=10 low-dose infected, n=10 high-dose infected) and C3HeB/FeJ mice infected with the H37Rv *Mtb* strain (n=5 uninfected, n=4 low-dose infected, n=5 high-dose infected) or the HN878 *Mtb* strain isolated (n=10 uninfected, n=9 low-dose infected, n=10 high-dose infected). As with the human data, we aggregated these cohorts into one mouse dataset with total n=88. The cohorts are described schematically in **Figure 1A-B**.

### Processing of Human Blood Sequencing Data

The human FASTQ raw blood sequencing TB data were obtained from GEO under accession numbers GSE107991 (London cohort) and GSE107992 (South Africa cohort). STAR (version 2.7.1a) was used to align the reads to the Hg38 human reference genome after which RSEM (version 1.3.1) was used to generate BAM files with gene and isoform expression level estimates from paired-end reads. The R package Tximport (version 1.16.1) was then used to generate count matrices from the gene expression level outputs from RSEM. Differential expression analysis was then performed using DESeq2 (version 1.28.1) with Benjamini-Hochberg FDR correction. Differentially expressed genes (DEGs) were identified using a significance threshold of *FDR<0.1* and *abs(log2FoldChange)>0.5* to identify differentially expressed genes (DEGs) between latently and actively infected TB patients in either the London or South Africa cohorts (**Figure S2**). DEGs were pooled across cohorts to create a single list of DEGs used to filter the human count matrices. The DESeq2 variance-stabilizing transformation was applied to the count matrices for each geographic cohort separately, followed by z-score normalization. The normalized and filtered count matrices were then aggregated to generate a single human dataset. This aggregated human dataset was filtered for one-to-one mouse-human gene homologs identified by the Bioconductor tools biomaRt and homologene (versions 2.44.0 and 1.4.68.19.3.27). Genes where more than one gene paired to a single gene of the opposite species were removed.

### Processing of Mouse Sequencing Data

Mouse FASTQ raw blood sequencing TB data was obtained from GEO under accession number GSE137092 and processed similarly to the human data, performing STAR alignment to the Mm10 mouse reference genome. Likewise, we then executed RSEM generation of gene and isoform expression level estimates from paired-end reads and Tximport generation of one count matrix for the whole mouse study cohort. Then, DESeq2 variance-stabilizing transformation and z-score normalization was applied to the count matrix. The normalized dataset was filtered for mouse orthologs of the human DEGs similar to what was done for the human dataset (**Figure S2**).

### Univariate comparisons of gene expression across species

Median gene expression of LTBI and ATB human samples were compared to the C57BL/6J and C3HeB/FeJ mouse cohorts based on a previously published gene set for discerning latent versus active TB cases in humans (**Figure 1C, Figure S1**). As mentioned previously, Bioconductor tools biomaRt and homologene (versions 2.44.0 and 1.4.68.19.3.27) were used to identify mouse homologs to the human gene sets. If the median gene expression was equal in both phenotypes of the same species, it was excluded from the alluvial plot, which was plotted using ggplot2 and ggalluvial (versions 3.4.2 and 0.12.5).

### Comparative enrichment of previously published TB gene signature

To compare the enrichment of previously published TB gene signatures in different phenotypic subsets of each species (**Figure 1D**), we utilized the *TBSignatureProfiler* package (version 1.8.0). We performed single sample GSEA using the package’s curated list of published TB signature gene sets containing 10 or more genes. Wilcoxon rank sum tests were performed on phenotype pairs (i.e., human ATB vs. LTBI; C3HeB/FeJ H37RV vs. HN878; C57/BL6 H37RV vs. HN878) to determine statistical significance.

### Regression of principal component scores onto covariates

Univariate regressions were performed to identify covariates that explain variation in score distributions on specific mouse principal components (**Figures 2C & S4B**). For binary covariates (mouse: mouse strain, Mtb strain; human: disease state, gender, cohort, BCG vaccination status), logistic regressions were performed and z-values were reported. For covariates that are continuous or 3+ ordinal categories (mouse: days since infection, bacterial dose; human: age), linear regressions were performed and t-values were reported. Benjamini-Hochberg multiple hypothesis correction was applied to the results for each species separately.

### Translatable Components Regression (TransComp-R)

Principal component analysis (PCA) was performed on the normalized mouse data using the stats package (version 3.6.2) to obtain the mouse sample scores and gene loadings on each mouse principal component (mPC). The first twenty mPCs explained 75% of variance in the mouse data and thus were kept for the next step of the analysis (**Figure S3A**). Human samples were projected into this twenty mPC space by multiplying the normalized human transcriptomic data matrix by the mouse loadings matrix, resulting in a matrix of human scores in the mPC space. For ease of comparison, the sign on mPC loadings was standardized such that positive values were associated with ATB. The variance of the human scores along all mPCs was totaled and used to normalize the amount of variance in the projected human data explained by each individual mPC. Down-selection of mPCs occurred in the following manner: (1) the first 7 mPCs were first selected as they explain 75% of the projected human variance and then (2) only mPCs with human ATB - LTBI effect size (including the full 95% confidence interval) above one were kept for the final logistic regression model. Univariate effect size, here parameterized by the Cohen’s d coefficient, was calculated for each mPC using the effsize R package (version 0.8.1). The projected human sample scores along these selected mPCs were then inputted into a logistic regression model using an 80:20 train-test split and 5-fold repeated cross validation with 10 repetitions. This model was trained for predicting human infection status (i.e., ATB vs LTBI) based on the human transcriptomic data projected into the mPCs.

### TransComp-R Model Validation

To test the likelihood of constructing a TransComp-R model with accuracy in predicting human TB outcomes similar to our final model, we compared our 5 mPC TransComp-R model to a series of null models with either 5 random mPCs or shuffled human phenotype labels. For the random mPC models, 5 mPCs between mPC1 and mPC88 were selected for a logistic regression model where projected human sample scores were regressed onto ATB and LTBI labels. This selection process of 5 random mPCs occurred 1000 times and the distribution of accuracy scores from 5-fold cross validation was plotted. For the null models with shuffled human phenotypic labels, the projected human samples on the five mPCs included in our final model (mPC1, mPC2, mPC3, mPC5, and mPC7) were regressed onto randomly shuffled ATB and LTBI labels. This was repeated 1000 times with newly shuffled labels each round and the distribution of accuracy scores from 5-fold cross validation was plotted. Both distributions of null models were statistically compared to the distribution of accuracy scores from the final TransComp-R logistic regression model using an 80:20 train-test split and 5-fold repeated cross validation with 10 repetitions.

### Comparison with mRNA signatures and gene sets

The loading scores for human genes on translatable component axes were compared with 67 previously-described human and mouse mRNA signatures using the single sample GSEA method in the “TBsignature profiler” package (12). Signatures that contained fewer than 10 genes in our dataset were excluded. Signature scores for each translatable component axis were compared by two way ANOVA, and clustered for presentation using the *hclust* package. GSEA analysis was performed using WebGestalt using the GSEA enrichment method, KEGG pathways between 5 and 2000 members. Significance was assessed using 1000 permutations.

### Bone-marrow derived macrophage production and culture conditions

C57BL/6J mice were purchased from Jackson Labs. Housing and experimentation were in accordance with the guidelines set forth by the Department of Animal Medicine of University of Massachusetts Medical School and Institutional Animal Care and Use Committee. BMDMs were generated from femur bone marrow via culture in DMEM containing 100 µM sodium pyruvate, 10 mM HEPES, 10% heat-inactivated FBS, and 20% conditioned media from L929 cells in non-treated tissue culture dishes for a total of 8 days. Additional media was added on day 3, and plates were washed with DPBS and received fresh media on day 5. Following differentiation, cells were harvested by rinsing plates with room temperature DPBS and then scraping the cells in cold DBPS. Collected cells were pelleted at 500xg for 5 min, resuspended in appropriate media, counted, and plated for treatments and infections. Following differentiation, cells were cultured in DMEM containing 100 µM sodium pyruvate, 10 mM HEPES, and 10% heat-inactivated FBS (cDMEM).

### UPR inhibition

For UPR inhibition studies, the following inhibitors and concentrations were used: 4µ8c (50 µM), Ceapin-A7 (10 µM), and GSK2606414 (1 µM). BMDMs were pre-treated with inhibitors or DMSO control for approximately 2 h prior to infection or tunicamycin treatment. For infection studies, pre-treatments were carried out at 1.5x desired concentration, subsequently adjusted to 1x upon addition of bacteria-containing media to wells. Inhibitors were maintained in media throughout the duration of the experiments.

### Mtb strains, culturing, and macrophage infection

Wildtype Mtb H37Rv was used for macrophage infection studies, with the exception of the bacterial uptake experiments which utilized H37Rv::pmCherry. Bacteria was cultured in Middlebrook 7h9 media with 0.2% glycerol, 0.05% Tween80 and 10% oleic acid-albumin-dextrose-catalase (OADC, BD). Prior to macrophage infections, bacteria were grown to mid-log phase (OD600 0.5-0.8). Bacteria was then collected, pelleted at 4000 RPM, resuspended in 30 ml cDMEM, pelleted at 4000 RPM, resuspended in 10 ml cDMEM, and spun at 700 RPM to pellet clumped bacteria. Supernatant was then transferred to a new tube and OD600 was measured to calculate MOIs for infection. Bacteria were added to cells at desired MOI (9 unless otherwise stated) and plates were spun at 500xg for 10 min. Cells were then returned to the incubator and infections allowed to proceed for 4 h. Cells were then washed to remove extracellular bacteria and fresh media containing appropriate inhibitors added. For RNAseq experiments, infections were allowed to proceed for 24 h before cells were lysed in TRIzol.

### RNA purification, transcriptional profiling, and quantitative PCR

For transcriptional profiling experiments, samples lysed in TRIzol (Thermo Fisher) were mixed with chloroform (5:1), incubated on shaker for at least 15 min, and were spun at 13,000 RPM at 4 for 15 min. Aqueous phase was then transferred to a fresh tube and an equal volume of 100% ethanol added. RNA was then isolated from solution using Direct-zol RNA Miniprep kit (Zymo Research) per manufacturer’s instructions. For each experimental condition, a total of 4 biological replicates were sequenced. Non-stranded library preparation with poly(A) enrichment and RNA sequencing was performed to recover >20,000,000 reads per sample. For qPCR experiments, cells were treated with tunicamycin (10 µM) ± UPR inhibition for 4 h before media was removed and cells were lysed with TRIzol. Lysed samples were added directly to Direct-zol RNA miniprep columns and isolated per manufacturer’s instructions. Reverse transcription was then performed using the iScript cDNA synthesis kit (Bio-Rad Laboratories). Quantitative PCR was performed with Luna Universal qPCR Master Mix (New England BioLabs) using an Applied Biosystems ViiA 7 Real-Time PCR system. Relative transcript abundance was calculated using the ΔΔCT method with target genes normalized against transcript levels of unspliced XBP1, for spliced XBP1, or β-actin, for HSPA5 and Trib3. The following primer sequences were used: spliced XBP1 F, 5’-ggtctgctgagtccgcagcagg-3’, R, 5’-gaaagggaggctggtaaggaac-3’ (41); unspliced XBP1 F, 5’-cagactatgtgcacctctgc-3’, R, 5’-cagggtccaacttgtccagaat-3’ (42); HSPA5 F, 5’-acttggggaccacctattcct-3’, R, 5’-atcgccaatcagacgctcc-3’ (43); Trib3 F, 5’-gcaaagcggctgatgtctg-3’, R, 5’-agagtcgtggaatgggtatctg-3’ (44–46); β-actin F, 5’-cgcagccactgtcgagtc-3’, R, 5’-ccttctgacccattcccacc-3’ (47).

### Mtb internalization

BMDMs were plated to 96-well plates and infected with H37Rv::pmCherry at indicated MOIs alongside UPR inhibitors or vehicle control (detailed above). Infection was allowed to proceed for 4 h before cells were washed with DPBS and fresh DPBS added to wells. Fluorescence was then measured using a Synergy H4 Hybrid Microplate Reader (BioTek). To account for background fluorescence, the mean signal from untreated, uninfected wells was subtracted from the other readings.

### RNAseq analysis

Sequencing file quality was verified using FastQC (48) version 0.11.5. Reads were trimmed using Cutadapt (49) version 4.1, then mapped to the mouse Ensembl GRCm38 genome (50) using STAR(51) version 2.5.2b. Transcripts were quantified using RSEM(52) version 1.3.3. Subsequent analysis was completed in R version 4.1.2. Files were imported using the Bioconductor package tximport (53) version 1.22.0. Libraries with a TPM less than or equal to 0.5 were excluded, and genes with fewer than 15 counts across all samples were excluded. Differential expression analysis was performed using the Bioconductor package DESeq2 (54) version 1.34.0. Pairwise comparisons between conditions were made using the Wald test. K-means clustering was performed on scaled, averaged normalized counts from 4 biological replicates/condition using Morpheus (55).

### Ingenuity Pathway Analysis and presentation

Pathway analyses was performed using Ingenuity Pathway Analysis (IPA) (Qiagen) software. For all analyses, pathways with a -log(p-value) > 1.3 and a z-score > 2 or <-2 were considered significantly enriched. TCs were analyzed using the Core Analysis function with a stringent filter limiting relationships to those occurring in immune cells. Major TC-associated pathways were identified as the 2 pathways with the highest and lowest z-scores, along with a significant p-value, for each TC. For K-means clustered data, IPA core analysis was used with default settings to discern the pathways most associated with each group. For associating BMDM RNAseq data with major TC pathways, core analysis was performed on genes showing significant (p-adj > 0.05) differential expression relative to the infected, uninhibited condition for each experimental group. Analysis was performed with a stringent filter limiting output to relationships occurring in murine cells.

### Programming

All analyses were done in R. R version 4.2.1 was used for preprocessing of the previously published human and mouse model RNASeq data and the TransComp-R modeling. R version 4.1.2 was used for BMDM RNASeq analyses.

### Study Approval

Animal housing and study protocols were in accordance with the guidelines set forth by the Department of Animal Medicine of Umass Chan Medical School and Institutional Animal Care and Use Committee.

## Data Availability

The publicly available data utilized in this study can be found on NCBI GEO under the following accession numbers: GSE107991 ( human London cohort), GSE107992 (human South Africa cohort) and GSE137092 (mouse cohort). The BMDM RNASeq results published in this study can be found on NCBI GEO under the follow accession number: XXXXX. All original code written for this study can be found at https://github.com/Lauffenburger-Lab.

## Funding Sources

This work was supported by NIH contract 75N93019C00071 to C.M.S. and D.A.L., and the Army Institute for Collaborative Biotechnologies collaborative agreement W911NF-19-2-0026 to D.A.L. Work by K.M.P. was supported in part by the National Science Foundation Graduate Research Fellowship (grant no. 1745302), the MIT-Takeda Fellowship Program, and Siebel Scholars. R.F was supported by NRSA fellowship AI176787.

## Acknowledgements

The authors would like to gratefully acknowledge members of the Lauffenburger lab, Sassetti Lab, and HI-IMPAcTB consortium for thoughtful feedback throughout the development of this project. The MIT co-authors would also like to thank Brian Joughin, Douglas Brubaker, and Meelim Lee for input on the modeling approach, along with Audrey Vargas, Elizabeth Chung, and Audrey Creighton for their related efforts.

## Supplementary Figures

**Figure S1.**
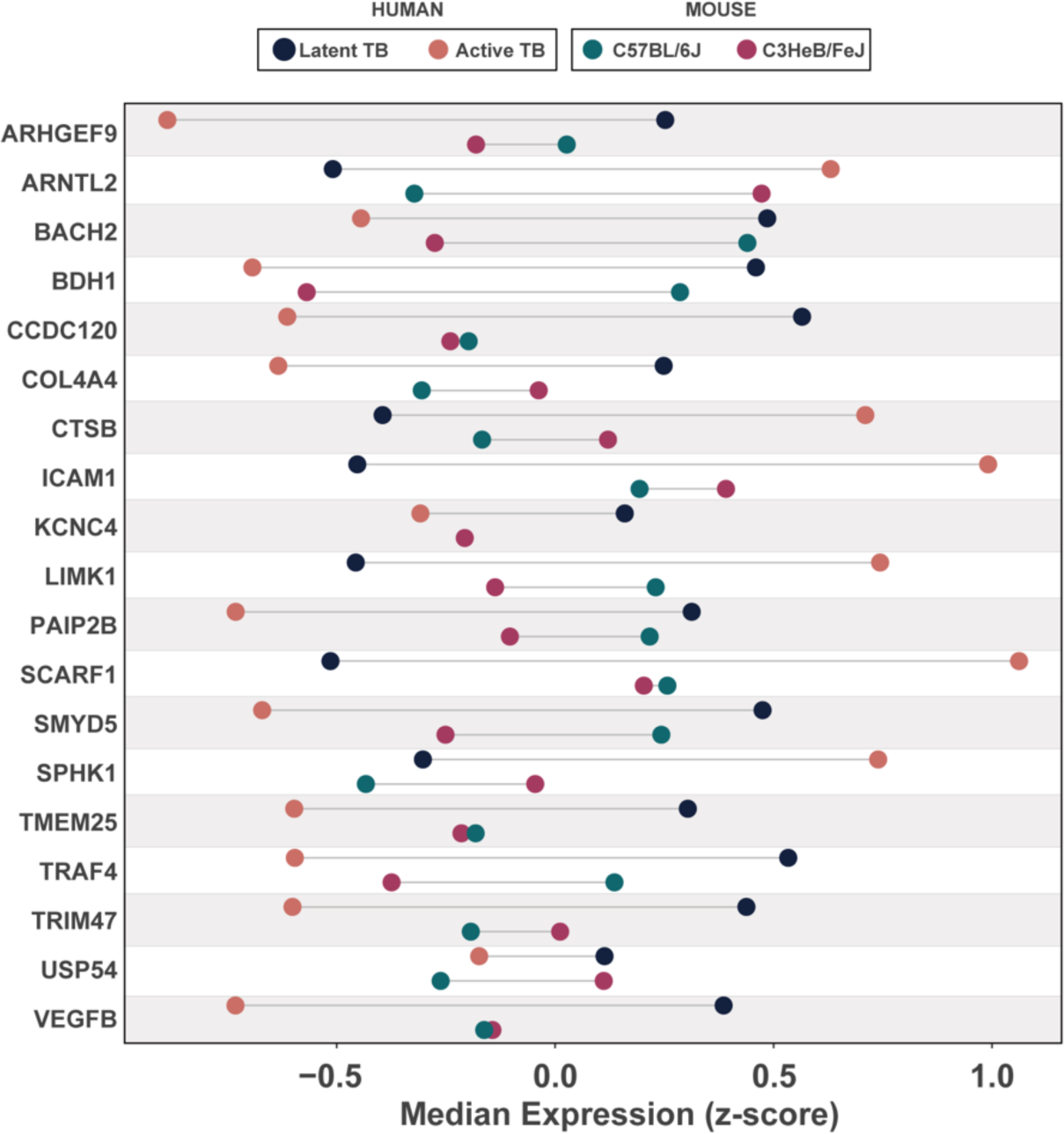
Gene expression of TB-associated genes differs between species. (A) Lollipop plot comparing the range of median gene expression z-scores in humans and mice for the 20 genes proposed by Singhania et al. (2018) to discriminate between active and latent tuberculosis infection in humans. APOL4 is not shown because there is no known mouse homolog.

**Figure S2.**
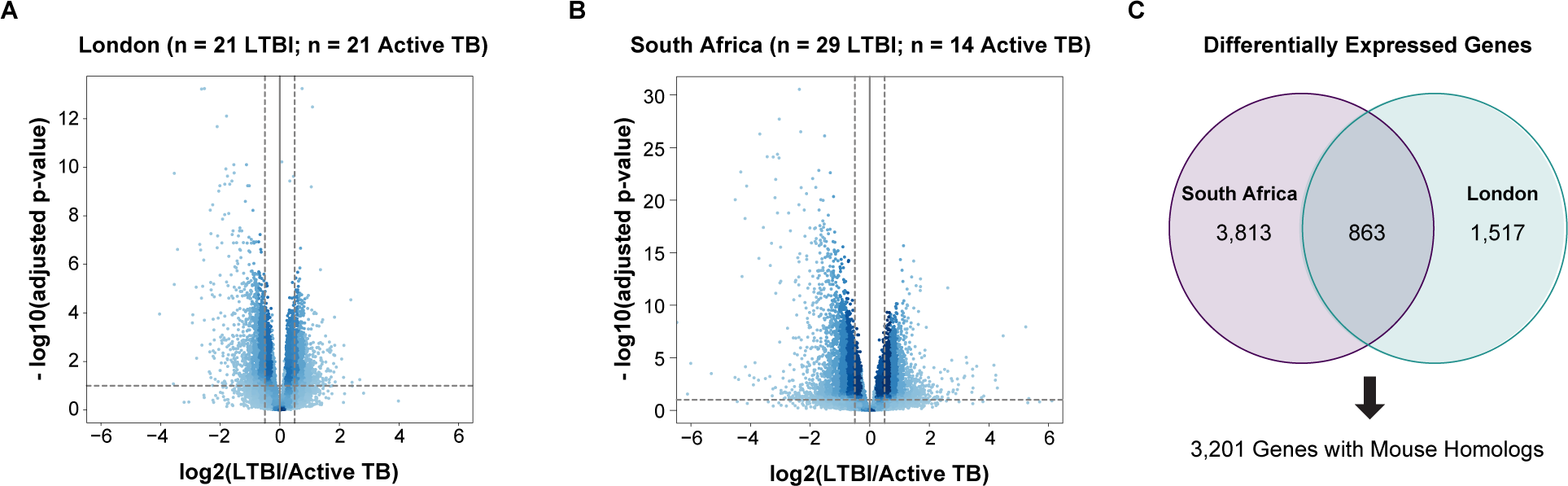
Identification of differentially expressed genes in each human cohort. Volcano plots of differentially expressed genes in London (A), and South Africa (B) cohorts. (C) Venn diagram illustrating genes differentially expressed similarly and differently across human cohorts.

**Figure S3.**
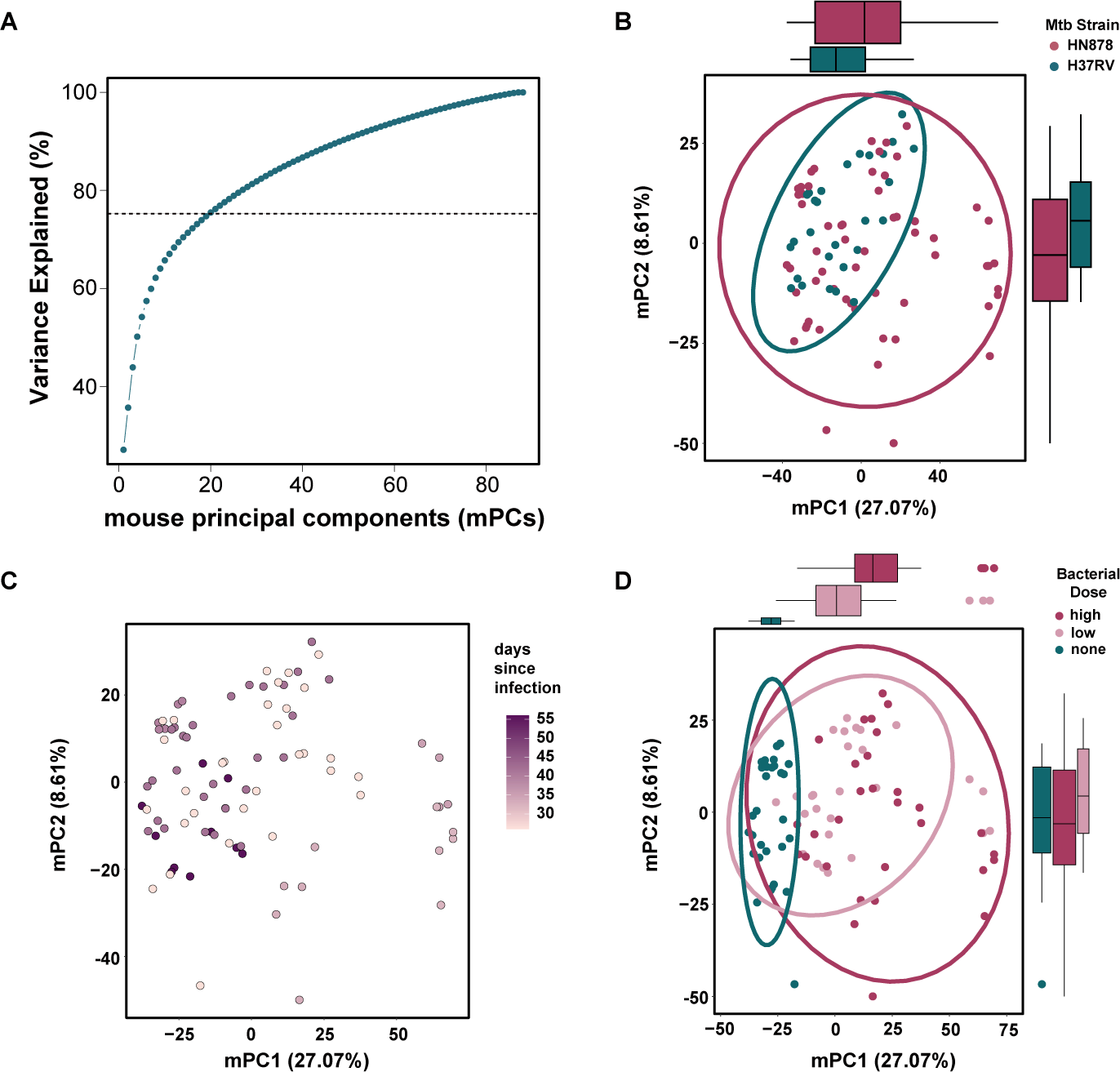
Transcriptional variation captured by the mouse PCA model. (A) Plot of variance explained across mouse principal components for the mouse PCA analysis. The first 20 PCs explain 75% of the variance and, thus, were kept for the TransComp-R model. (B) – (D) Principal component analysis on mouse homologs of the human differentially expressed genes in latent vs. active tuberculosis. Each data point represents a mouse sample in the PC1 vs. PC2 space. Data points are colored by (B) Mtb strain, (C) days since infection and (D) bacterial dose. The colored ellipses in (B) and (D) represent the 95% confidence interval for each phenotype’s distribution in the PC1 vs. PC2 space. Similarly, boxplots on the x- and y-axes in (B) and (D) illustrate the distributions of each phenotype.

**Figure S4.**
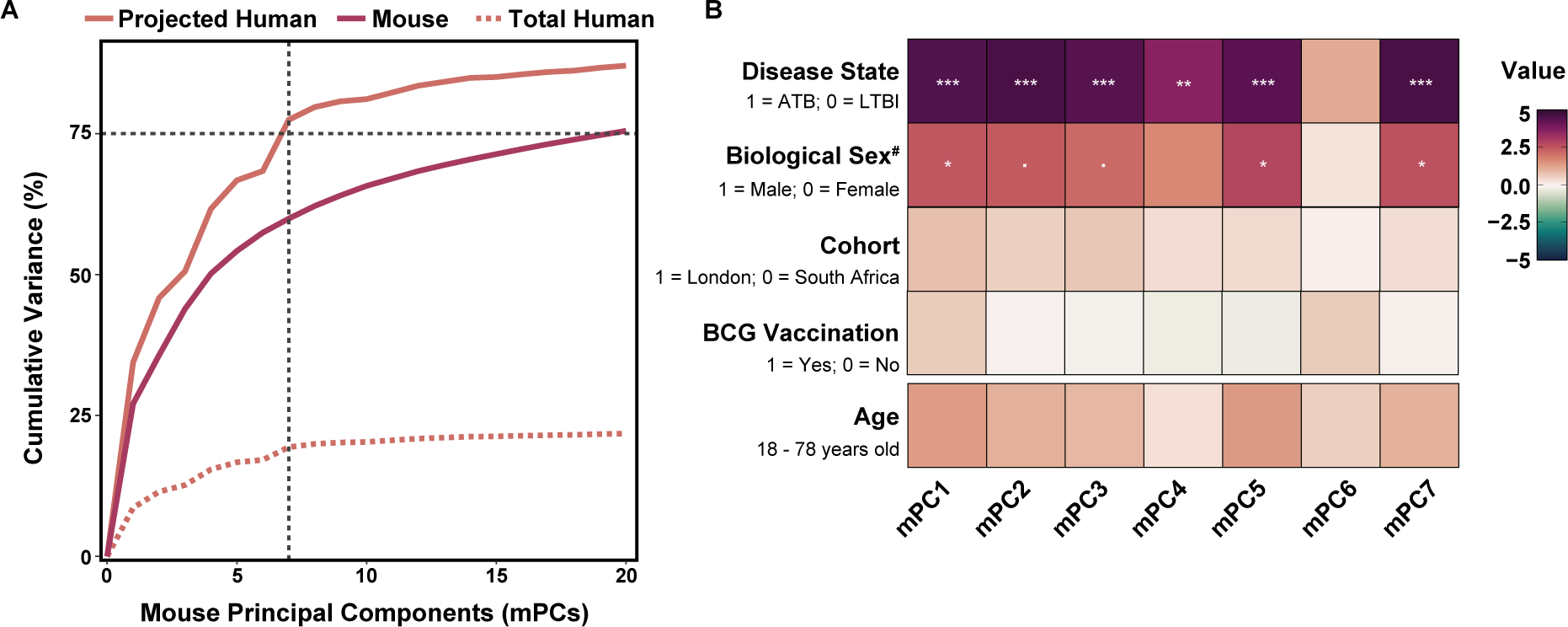
Human transcriptional variation captured by the mouse principal components. (A) Cumulative variance explained by the mPCs for both the human and mouse data. Dashed black lines indicate where 75% of the projected human variance is explained. The dashed peach line represents the total raw human variance explained by the mPCs. (B) Regression models where projected human sample scores in the mouse principal component (mPC) space were regressed on human disease state, reported gender, geographic cohort, BCG vaccination status, and age. Z-values are shown for the logistic regression models (gender, cohort, BCG vaccination). T-values are shown for the linear regression models (age). Positive values (magenta) correspond with the ATB phenotype, Male patients, the London cohort, and previous BCG vaccination. ‘.’ indicates p<0.1, * indicates p<0.05, ** indicates p<0.01 and *** indicates p<0.001 after Benjamini-Hochberg multiple hypothesis correction. ^#^Here we label our metadata feature as biological sex as patients reported either “male” or “female.” This feature was labeled as “gender” in the original study published by Berry et al. (40).It is unclear if patients were asked to report biological sex or gender. If there was no gender reported for an individual, the patient was not included in the regression model.

**Figure S5.**
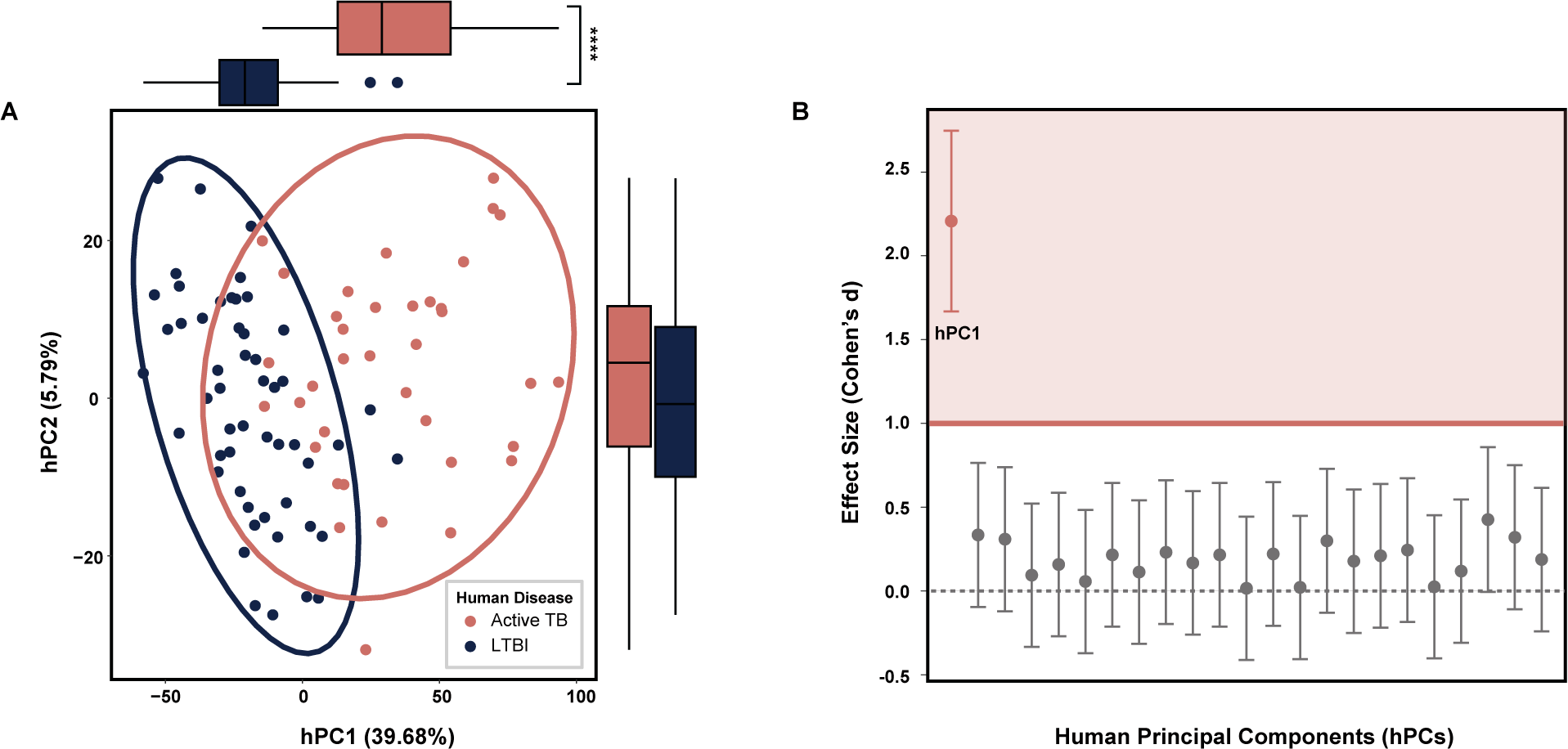
PCA analysis performed directly on human transcriptomics data captures most disease phenotype-related variation on the first principal component. (A) Principal component analysis on human differentially expressed genes in latent vs. active tuberculosis. Each data point represents a human sample, and the colored ellipses represent the 95% confidence interval for each strain’s distribution in the PC1 vs. PC2 space. Boxplots on the x- and y-axes illustrate the distribution of the active tuberculosis (salmon) and LTBI (navy) samples on each axis respectively. Wilcoxon Rank Sum Tests were performed on the scores of each mouse strain for each principal component. **** p-value < 0.0001. (B) Plot comparing the univariate effect size observed between the ATB and LTBI human samples projected into each individual human PC (hPC). Effect sizes, measured by Cohen’s d coefficient, with a magnitude greater than one are shaded pink. Confidence intervals for each Cohen’s d coefficient are indicated by error bars.

**Figure S6.**
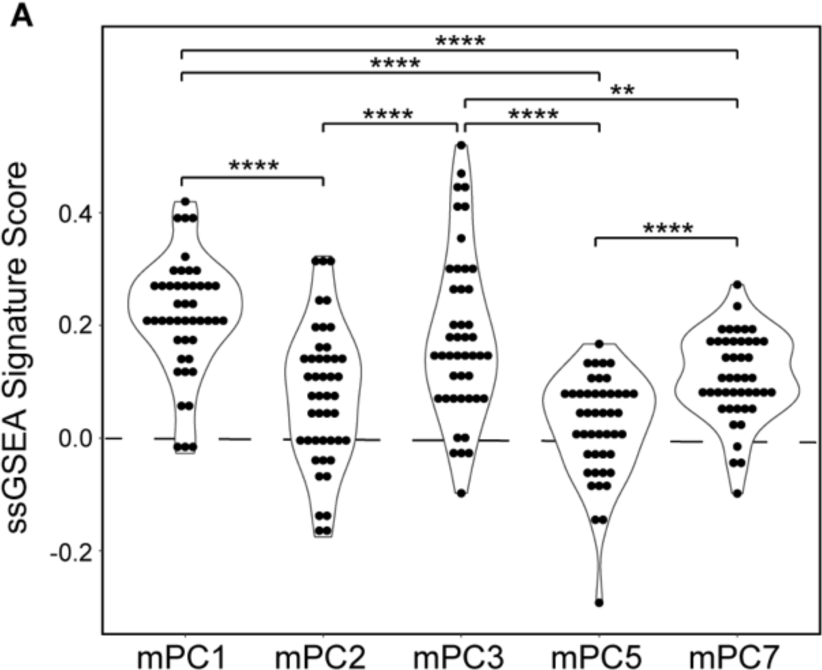
Enrichment of previously published gene signatures distinguishing ATB and LTBI on each translatable component. (A) ssGSEA was performed using the “TBsignature profiler” package (12) in R. Signatures that contained fewer than 10 genes in our dataset were excluded. Pairwise paired sample Wilcoxon tests were performed to determine statistical significance and p-values were adjusted using Benjamini-Hochberg multiple hypothesis correction. **** p<0.0001; *** p<0.001; **p<0.01; *p<0.05

**Figure S7.**
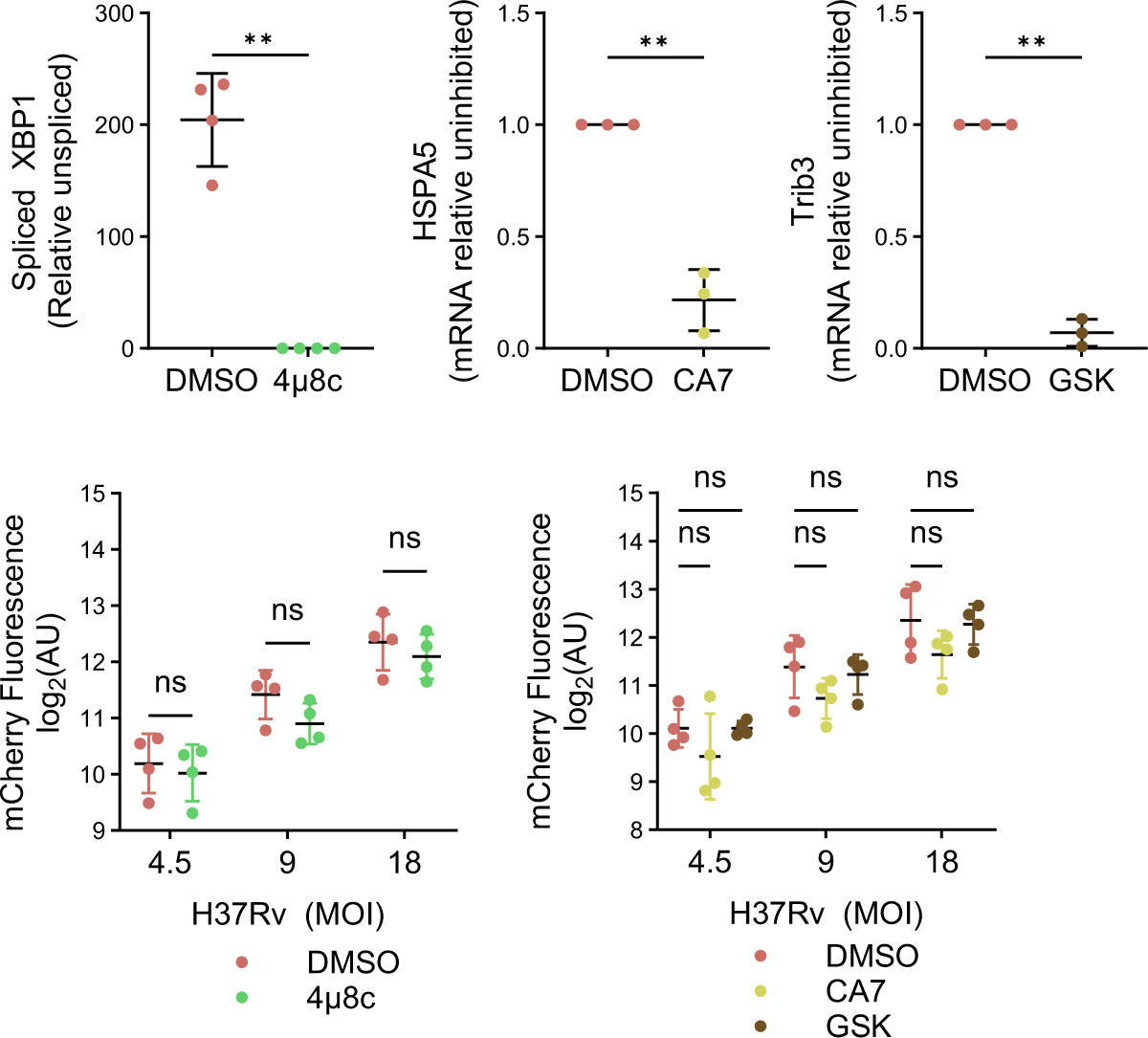
UPR inhibitors antagonize the appropriate pathway and do not inhibit Mtb internalization. (A) BMDMs were treated with tunicamycin in the presence of UPR inhibitors or vehicle control and transcript levels of UPR targets were determined via qPCR. For spliced XBP1, data is normalized to non-tunicamycin treated control. Bars represent mean ± SD of 4 biological replicates. Significance was determined using Welch’s t-test; ** indicates P < 0.01. (B,C) BMDMs were infected with mCherry-expressing H37Rv at the indicated MOIs. 4 h post-infection, mCherry fluorescence was measured as a proxy for bacteria uptake. Bars represent mean ± SD of 4 biological replicates. Significance was determined by 2way ANOVA with Sidak’s (B) or Tukey’s (C) multiple comparison testing; “ns” indicates P > 0.05.

## Notes

### Competing Interest Statement

The authors have declared no competing interest.

